# How opsins diversified after the teleost whole-genome duplication: Insights from two parietopsins of the red piranha, *Pygocentrus nattereri*

**DOI:** 10.1101/2025.09.17.675483

**Authors:** Keita Sato, Takahiro Yamashita, Kazumi Sakai, Hideyo Ohuchi, Takehiro G. Kusakabe, Fuki Gyoja

## Abstract

**Background:** The teleost whole-genome duplication (TGD) contributed to functional diversification of opsins. Some TGD paralogs, including those of parapinopsin (PP), Vertebrate Ancient (VA) opsin, and long wavelength-sensitive opsin (LWS), show different absorption spectra and/or expression patterns. However, our knowledge of detailed evolutionary processes and mechanisms by which TGD contributed to opsin diversification is still limited.

**Results:** Here, we report that several characins, including the red piranha (*Pygocentrus nattereri*), have retained TGD paralogs of parietopsin (PT1 and PT2). While most teleosts, including zebrafish, have only PT1, the Mexican tetra (*Astyanax mexicanus*) and catfishes have only PT2. To assess the degree of functional diversification between PT1 and PT2, we characterized spectral properties, intracellular signaling properties, and expression patterns. Maximum absorption spectra differ slightly among PTs. Those of red piranha PT1, PT2, Mexican tetra PT2, and Japanese catfish (*Silurus asotus*) PT2 are located at 517 nm, 528 nm, 517 nm, and ∼535 nm, respectively. In a heterologous cell-based assay, red piranha PT1 increased cellular cAMP in response to light, whereas PT2 decreased it, while neither induced a clear Ca²⁺ response under the conditions tested. Fluorescence *in situ* hybridization showed that (1) piranha *PT1* and *PT2* are expressed in the same pineal cells, and (2) they are also co-expressed with *PP1*.

**Conclusions:** This study illustrates the importance of characterizing intracellular signaling properties to understand how TGD contributed to functional diversification of opsins.

## Background

Opsins are light-sensitive, G-protein-coupled receptors (GPCRs) found in animals. Traditionally, they have been categorized into seven to nine superfamilies [1,2]. For most of them, the Urbilateria are presumed to have had at least one member [3]. This repertoire has been further expanded in both protostomes and deuterostomes. How and why such diversity was achieved, and how it enabled sophisticated animal photoreception, are important questions.

The teleost whole-genome duplication (TGD) contributed greatly to opsin diversification (Fig. 1A) [4]. For example, parapinopsins (PPs) 1 and 2 are TGD paralogs [5]. Many extant teleosts have both. While PP1s are UV-sensitive, like lamprey PP, PP2s are visible-light sensitive [5]. PP1 and PP2 are expressed in a complementary fashion in zebrafish pineal gland, and PP1 is co-expressed with parietopsin (PT) [5].

**Figure 1.**
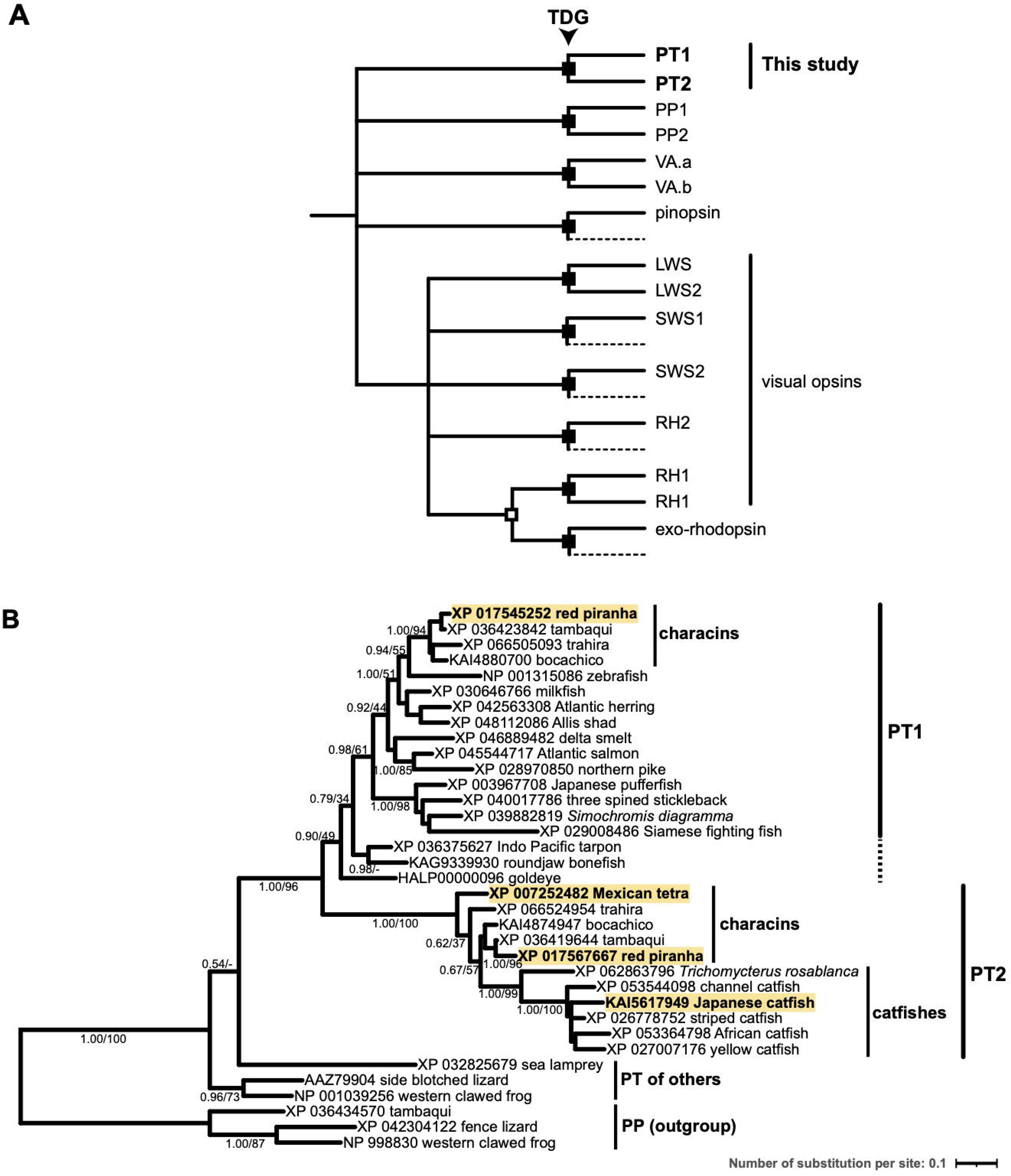
The teleost whole-genome duplication (TGD) contributed greatly to opsin diversification. (A) A schematic representation of vertebrate visual and non-visual opsin diversification in fishes, based on [5–7,9–12,26,28,44]. Gene duplications by TGD are shown with closed squares. RH1 experienced retrotransposition/retroduplication prior to TGD (shown with an open square). (B) Teleost PTs form two distinct clusters, PT1 and PT2. Most species have only PT1. Several characins, including the red piranha, have both PT1 and PT2. The Mexican tetra and catfishes have only PT2. Although PTs of early-branching teleosts tend to be clustered with PT1, fidelity is not necessarily high. This tree was generated using the Bayesian Inference (BI) method. The maximum-likelihood (ML) method yielded a similar result. Posterior probabilities of BI and bootstrap values of ML are shown in this order at major nodes. PTs used for spectroscopic measurements (Fig. 3) are highlighted. Amino acid residues that are conserved in either PT1 or PT2 are shown in Supplementary Fig. S8. Alignments are available in Supplementary Data 1. See [45] for the goldeye PT. Our recent extensive survey failed to find PTs in non-teleost fishes, such as the spotted gar, the Siberian sturgeon, and the gray bichir [17]. Abbreviations: PT, parietopsin; PP, parapinopsin; VA, vertebrate ancient opsin; LWS, long wavelength-sensitive opsin; SWS1/2, short wavelength-sensitive opsins; RH2, rhodopsin-like 2; RH1, rhodopsin.

TGD paralogs of long wavelength-sensitive opsin (LWS) are another example. While one paralog remains red-light sensitive, the other (LWS2) acquired green-light sensitivity [6]. Unlike PPs, LWS2s are reported only from osteoglossomorphs and characins, suggesting secondary losses in many lineages [6]. TGD paralogs have also been reported in other opsin subfamilies, including RH1, VA opsin, and Opn4/melanopsin [7–11]. Unlike PP and LWS paralogs, spectral properties and/or expression patterns between these paralogs are comparable [7,12]. For example, the zebrafish has TGD paralogs of VA opsin, VA.a and VA.b. They are expressed mutually exclusively in the larval eye and brain [7]. However, their spectral properties are comparable. The absorption maximum of the former is located at 510 nm, and the latter at 505 nm [7]. Therefore, diversification after TGD may differ among opsins.

PT is a green-sensitive opsin first isolated from lizard parietal eyes [13]. Parietal eyes may be involved in detection of dawn and dusk, sky polarization pattern, and magnetic field, although underlying molecular mechanisms remain to be addressed [14–16].

Recently, we reported that a few characins, including the red piranha (*Pygocentrus nattereri*), have two PT paralogs [17]. Here, we report that they are TGD paralogs (PT1 and PT2). To our knowledge, this is the first report that TGD paralogs of PT are retained. Spectral properties of PT1s and PT2s are comparable. In a heterologous cell-based assay, red piranha PT1 increased cellular cAMP in response to light, whereas PT2 decreased it, while neither induced a clear Ca²⁺ response under the conditions tested. We also surveyed mRNA distributions of *PT1* and *PT2* in the red piranha. We found that *PT1* and *PT2*, together with *PP1*, are co-expressed in pineal cells. Based on these discoveries, we discuss how functional diversification of opsins may have occurred after gene duplications.

## Results

### PT1 and PT2 are TGD paralogs

Recently, we completed a high-quality catalog of vertebrate non-visual opsins and reported that while most teleosts possess one parietopsin (PT) gene, a few characins, including the red piranha (*P. nattereri*), have two paralogs [17]. We speculated that they are TGD paralogs since (1) they were unlikely to be recently tandem-duplicated, judging from their location on different chromosomes, and (2) they were also unlikely to be produced by retrotransposition/retroduplication because both have three introns. To further study this, we performed molecular phylogenetic analyses. Teleost PTs form two distinct clusters, PT1 and PT2 (Fig. 1B). The two PTs of characins were assigned to PT1 and PT2, respectively (Fig. 1B). While most extant teleosts only have PT1, the Mexican tetra (*A. mexicanus*) and catfishes only have PT2 (Fig. 1B).

Although this suggests that PT1 and PT2 are TGD paralogs, we found few, if any other TGD paralog candidates around these loci (Fig. 2A). Because microsyntenic comparison with the pre-duplicated state is useful to infer evolutionary relationships between paralogs [18,19], we manually traced microsyntenic transitions. We deduced the teleost ancestral state (Fig. 2B) using three Elopomorpha species, the Indo-pacific tarpon (*Megalops cyprinoides*), the roundjaw bonefish (*Albula glossodonta*), and the European eel (*Anguilla anguilla*), and two non-teleosts, the spotted gar (*Lepisosteus oculatus*), and the gray bichir (*Polypterus senegalus*) (Fig. 2C-G). Comparing the teleost ancestral state (Fig. 2B) with genomic regions around *PT1* and *PT2* (Fig. 2A), we concluded that genes surrounding them are likely descendants of genes around the PT locus in the pre-duplication state.

**Figure 2.**
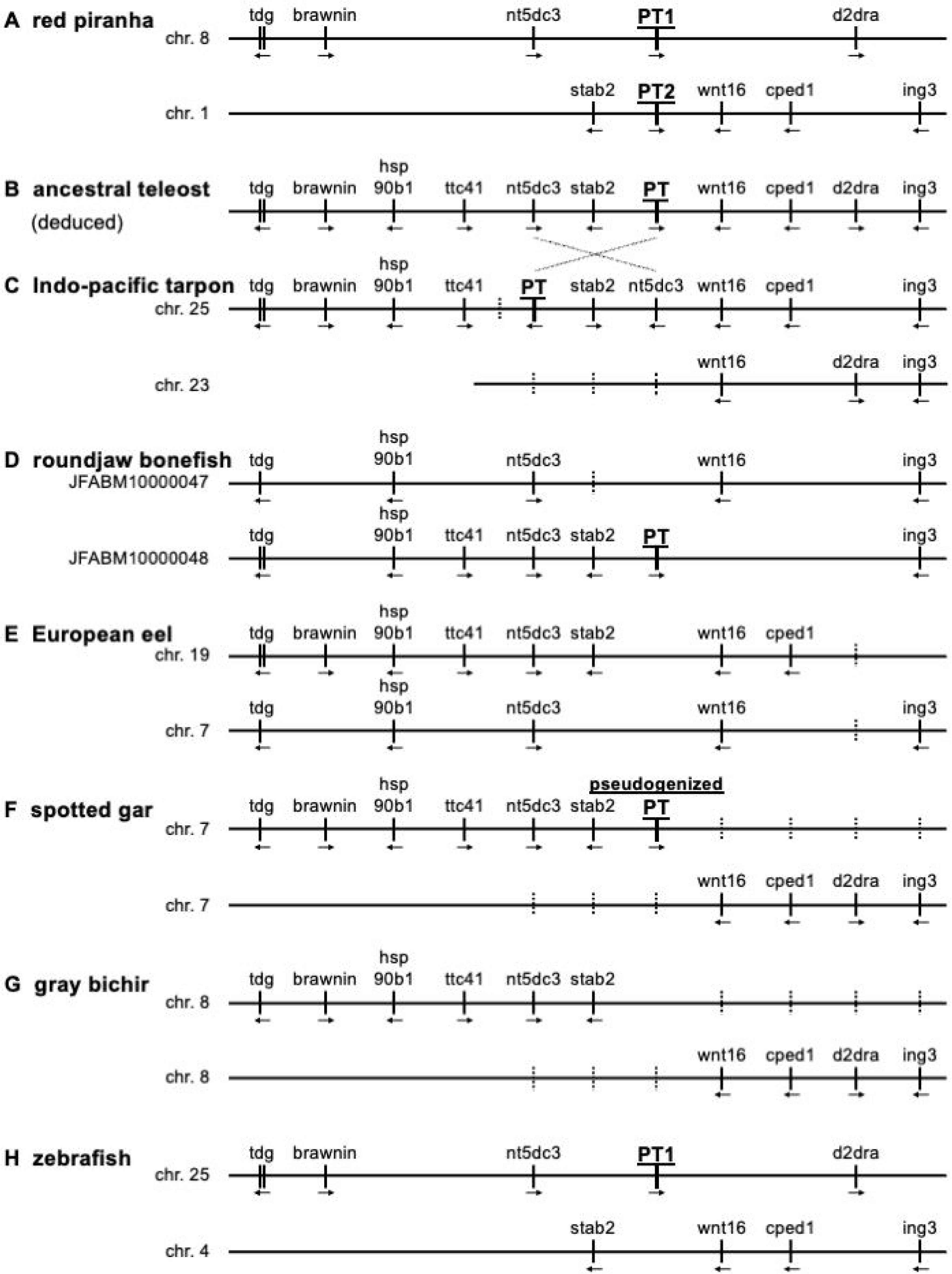
PT1 and PT2 are TGD paralogs. (A) Microsynteny around red piranha PT1 and PT2. Few TGD paralog candidates other than PTs are observed. (B) Ancestral microsynteny just before TGD was deduced from that of (C) the Indo-pacific tarpon, (D) the roundjaw bonefish, (E) the European eel, (F) the spotted gar, and (G) the gray bichir. Small-scale inversion has probably occurred in the tarpon lineage. An intrachromosomal rearrangement has probably occurred in the ancestral teleost lineage after its separation from the gar lineage. We could not assign PTs of the tarpon and the bonefish to either PT1 or PT2. This might be due to "delayed diploidization after TGD", recently proposed by [19]. (H) zebrafish corresponding genomic regions, which were used for deduction of the ancestral chromosomal assignment. Predicted protein-coding genes, other than those used for microsyntenic analyses, are shown with dashed lines. GenBank/DDBJ/EMBL accession IDs of proteins encoded by these genes are provided in Supplementary Table S1.

Recent studies have reported detailed macrosyntenic evolution of teleosts [19,20]. Parey et al. [19] deduced that zebrafish genomic regions corresponding to piranha microsyntenic blocks (Fig. 2H) descended from homeologous ancestral chromosomes, 11a (corresponding to *PT2*) and 11b (*PT1*), respectively. Considering these matters, we concluded that PT1 and PT2 are TGD paralogs.

### Spectral properties of PTs of red piranha, Mexican tetra, and Japanese catfish

Since absorption spectra sometimes differ among TGD opsin paralogs [5,6], we next investigated whether this is also true of PT. We analyzed absorption spectra of PT1 and PT2 of the red piranha. We also examined PT2s of the Mexican tetra and the Japanese catfish, which have lost their PT1s secondarily. Recombinant pigments were expressed in HEK293S cells, reconstituted with 11-*cis* retinal, solubilized in 1% n-dodecyl-β-D-maltoside, and photobleached in the presence of 100 mM NH_2_OH to convert all photointermediates to retinaloxime. Subtracting absorption spectra recorded after photobleaching from those before photobleaching produced difference spectra with a peak in the visible region corresponding to the absorption maximum of the 11-*cis* retinal-bound pigment (Fig. 3; Supplementary Fig. S1). Absorption maxima of these difference spectra occurred at 517 nm for red piranha PT1, 528 nm for red piranha PT2, 517 nm for Mexican tetra PT2, and ∼535 nm for Japanese catfish PT2. Absorption maxima of PTs have been reported as follows: 520 nm for zebrafish PT1 [21], 522 nm for the side-blotched lizard [13], 520 nm for the western clawed frog [22]. We further compared residues surrounding the retinal chromophore. However, in this comparison, no single residue or simple motif clearly correlated with observed differences in absorption maxima among the PTs examined (Supplementary Fig. S2). Taken together, these results suggest that (1) spectral properties apparently differ little between PT1 and PT2, and (2) spectral properties are slightly divergent among PTs, especially among PT2s.

**Figure 3.**
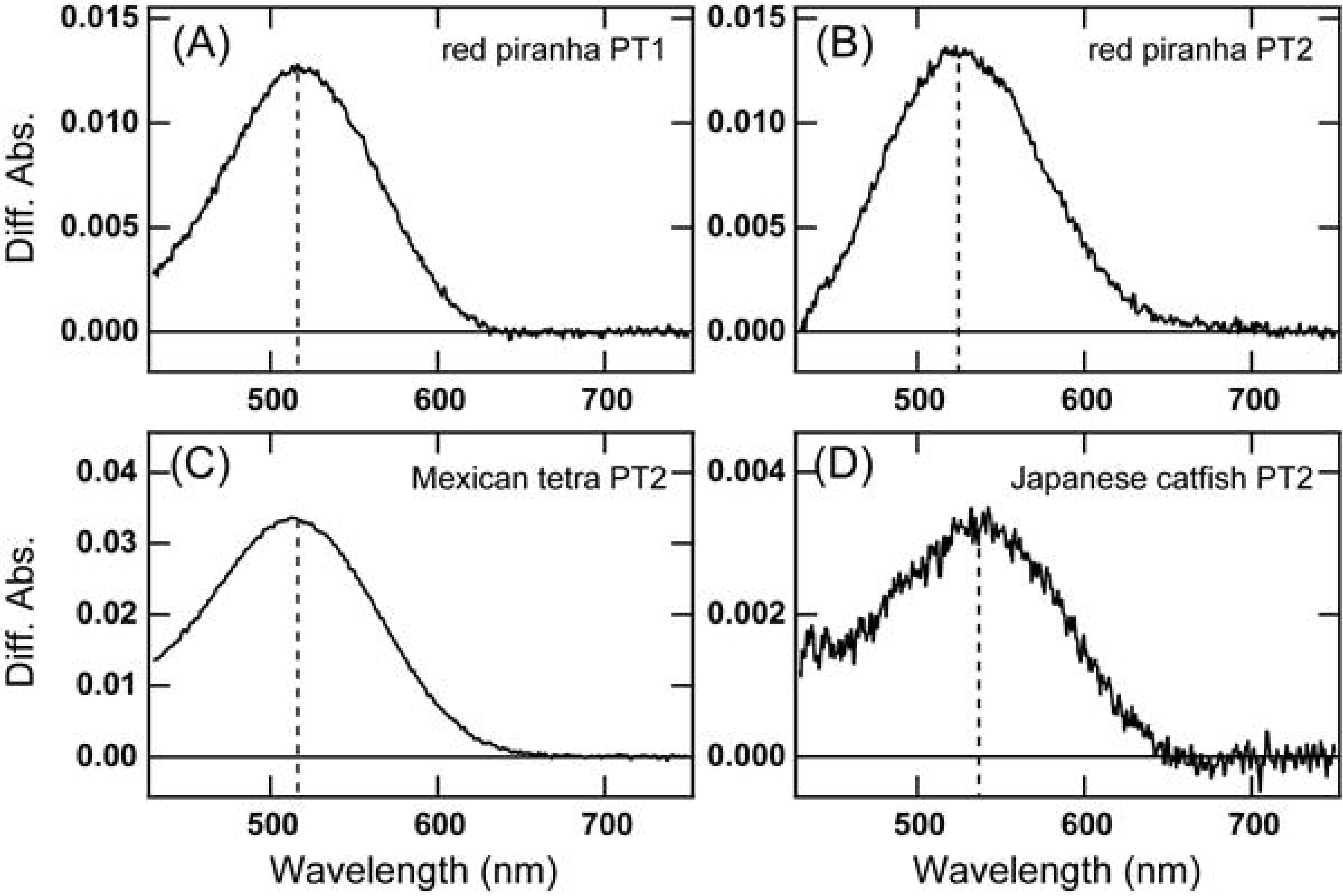
Absorption spectra of PT1 and PT2. Cell membranes expressing red piranha PT1 (A), red piranha PT2 (B), Mexican tetra PT2 (C) or Japanese catfish PT2 (D) after addition of 11-*cis* retinal were solubilized with 1% DDM and their absorption spectra were calculated by subtracting the spectrum after red light (>600 nm) irradiation, from that before irradiation in the presence of 100 mM NH_2_OH. Spectra, normalized to be ∼ 1.0 at the positive maximum, are compared in Supplementary Figure S1.

### Red piranha PT1 increases cellular cAMP, whereas PT2 decreases it in response to light

Next, to examine intracellular signaling properties of red piranha PT1 and PT2, we analyzed light-dependent changes in cAMP levels using the GloSensor 22F assay in cultured cells. Pigments were reconstituted with 11-*cis* retinal prior to the assay to regenerate their dark state. Upon light irradiation, cells expressing PT1 showed an increase in luminescence, indicating that PT1 increases cellular cAMP levels in response to light (Fig. 4A). This light-dependent increase was also observed after forskolin was added during the recording to elevate intracellular cAMP levels (Supplementary Fig. S3). In contrast, when intracellular cAMP levels were elevated by addition of forskolin, light stimulation of PT2-expressing cells caused a decrease in luminescence, indicating that PT2 decreases cellular cAMP levels in response to light (Fig. 4B). We also tested whether PT1 or PT2 could induce intracellular Ca²⁺ responses using an aequorin luminescence-based assay. As a positive control, mouse Opn4 evoked a clear light-dependent transient elevation of the Ca²⁺ level (Fig. 4C). By contrast, PT1- and PT2-expressing cells showed no clear increase in the Ca²⁺ level (Fig. 4C, D). These results indicate that in this heterologous expression system using cultured cells, red piranha PT1 and PT2 primarily modulate intracellular cAMP levels rather than triggering detectable Ca²⁺ mobilization under the present experimental conditions.

**Figure 4.**
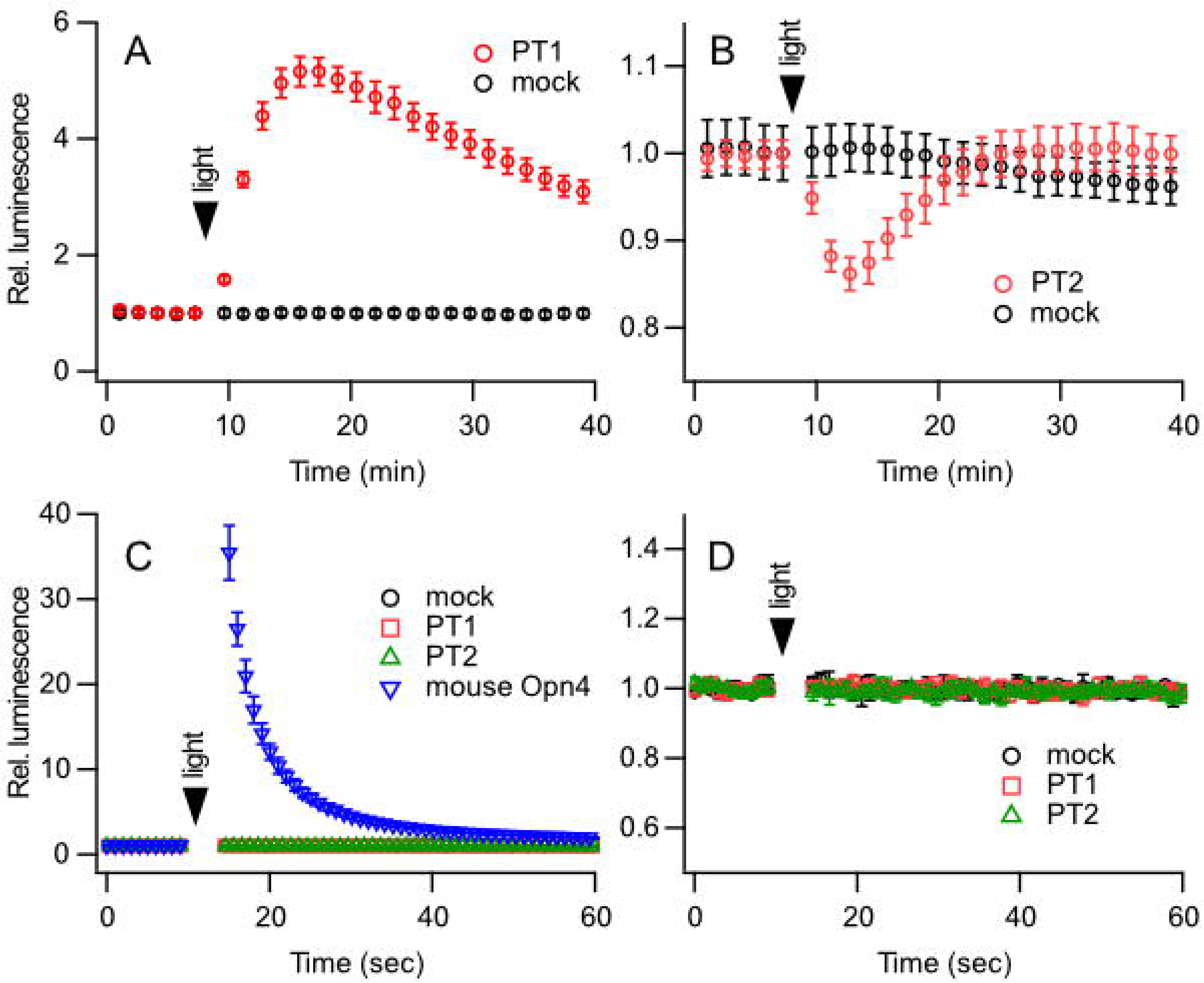
Cellular responses of red piranha PT1 and PT2 to light stimulation. (A, B) Time courses of relative luminescence in the GloSensor 22F cAMP assay in 293T cells expressing red piranha PT1 (A, red) or red piranha PT2 (B, red), compared with mock-transfected cells (black). A decrease of cAMP was assayed in the presence of 2 μM forskolin (B). (C) Time courses of relative aequorin luminescence in 293S cells for mock-transfected cells (black circles), PT1-expressing cells (red squares), PT2-expressing cells (green triangles), and mouse Opn4-expressing cells (blue inverted triangles). Mouse Opn4 was included as a positive control for the aequorin assay. (D) Enlarged view of panel C showing the traces for mock PT1 and PT2. No clear response was detected for PT1 or PT2 under these conditions. Data are shown as mean ± SEM (n = 3). Relative luminescence was normalized to the baseline before light stimulation.

### PT1, PT2, and PP1 are co-expressed in pineal cells of the red piranha

Some TGD opsin paralogs show different expression patterns [5,7]. To examine whether this is also true for PTs, we next investigated mRNA distributions of *PT1* and *PT2* in the red piranha. PT expression in teleosts has been reported almost exclusively in the pineal gland [21]. cDNA fragments of *PT1* and *PT2* were amplified from first-strand cDNA prepared from this tissue, but not from eyes. Therefore, we focused on the pineal gland, using both chromogenic and fluorescence *in situ* hybridization. Frontal brain sections containing the pineal gland were prepared. Hybridization signals for *PT1* and *PT2* mRNAs appeared in the dorsal region of the rostral pineal gland (Fig. 5A, B, E, F), but were absent from the caudal portion (Fig. 5C, D, G, H). This pattern parallels *PT* expression reported in zebrafish and river lamprey [21,23]. To determine whether the two *PT* genes are co-expressed in the same cells, we next performed fluorescence *in situ* hybridization on the same sections. We also examined the UV-sensitive parapinopsin *PP1*, because zebrafish and the green iguana show almost complete co-expression of *PP* and *PT*, whereas the river lamprey shows no overlap [21,23,24]. *PP1* signals were detected in the same region as *PT1* and *PT2* (Fig. 5I-M), and a substantial number of *PP1* positive cells appeared to co express *PT1* and *PT2* (Fig. 5N-R). Quantitative analysis of these images revealed a co-expression pattern with several distinct cell populations (Fig. 5S). Among all *PP1*-positive cells (n = 377), the largest group was that which co-expressed both *PT1* and *PT2* (43.5%, 164 cells), although substantial populations of cells expressed only *PP1* (31.0%, 117 cells) or co-expressed *PP1* with *PT2* (21.8%, 82 cells). Furthermore, this analysis highlighted strong co-expression between *PT1* and *PT2*. Of all *PT1*-expressing cells (n = 216), a vast majority (86.6%, 187 cells) also expressed *PT2*. In striking contrast, cells expressing only *PT1* were rare, constituting just 6.9% (15 cells) of this population. In addition to these expression patterns, we quantified transcript spot counts per cell and found that *PT1* levels were significantly lower than those of *PP1* and *PT2* (Fig. 5T). This trend was independently validated by RT-qPCR quantification using external RNA standards (Fig. 5U). These results indicate that the three opsins are often co-expressed, involving multiple cell populations with heterogeneous expression profiles.

**Figure 5.**
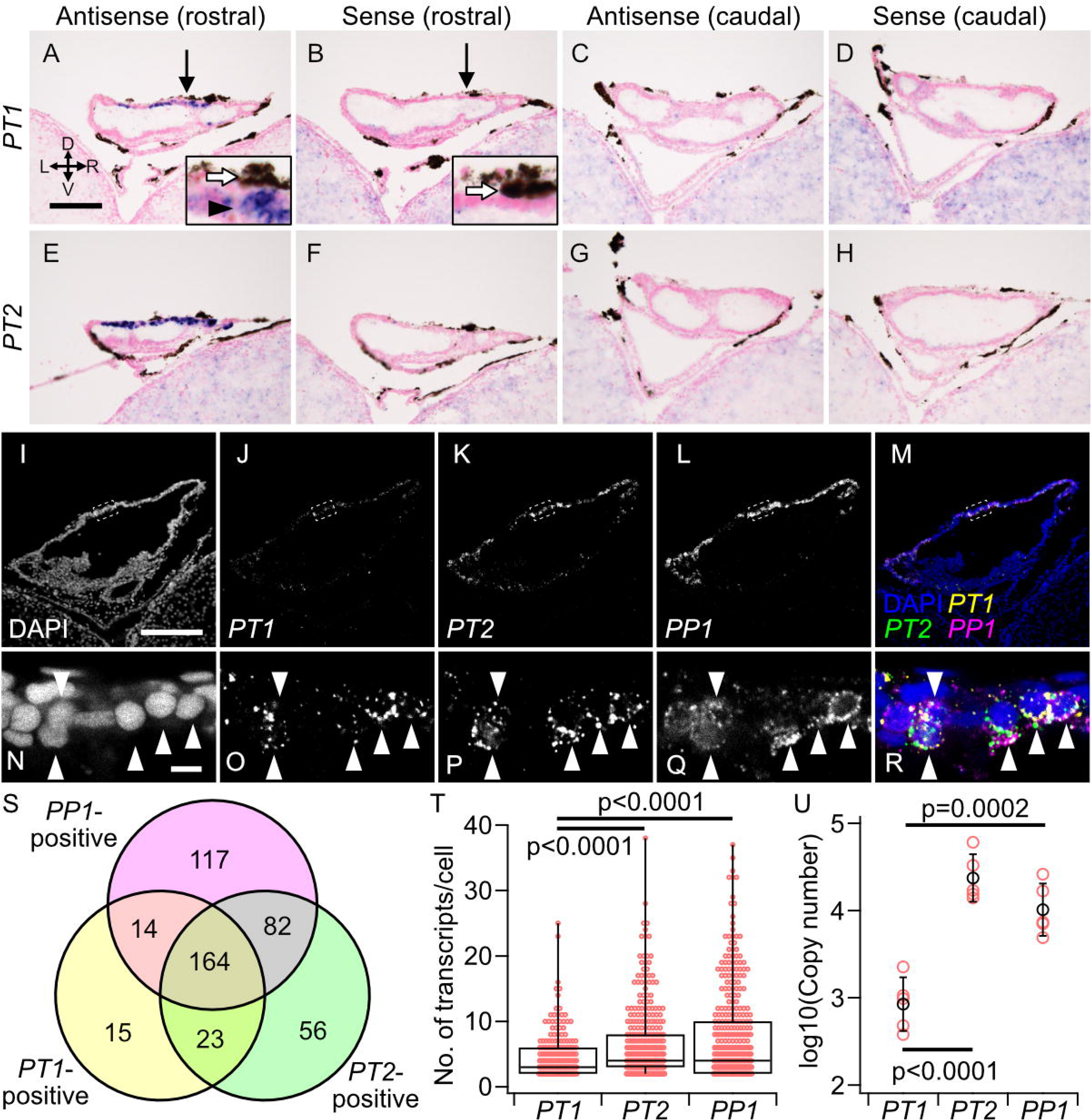
mRNA localization of red piranha parietopsins in the pineal gland. (A–H) Chromogenic *in situ* hybridization for *PT1* (A–D) and *PT2* (E-H) in the red piranha pineal gland. All images are frontal sections. Dorsal is up, and the left side of the animal is on the left (axes indicated). Panels A, C, E, and G show sections hybridized with antisense probes. Consecutive sections hybridized with sense probes are shown in panels B, D, F, and H, corresponding to panels A, C, E, and G, respectively. Panels A, B, E, and F are from the rostral half of the pineal gland, whereas panels C, D, G, and H are from the caudal half. Sections were counterstained with Nuclear Fast Red. Insets in panels A and B show an enlarged view of the region indicated by arrows. The blue–purple signal (arrowheads in insets) represents NBT/BCIP precipitate produced by alkaline phosphatase and indicates a specific hybridization signal. The brown-to-black granules visible in both antisense and sense panels (open arrows in insets) are endogenous melanin pigment. (I-R) Fluorescence *in situ* hybridization (FISH) for *PT1* (J, O), *PT2* (K, P), and *PP1* (L, Q). *PT1*, *PT2*, and *PP1* were visualized with ATTO 488, ATTO 565, and ATTO 633, respectively. Nuclei were stained with DAPI (I, N). Panels M and R show merged pseudo-color images of panels I-L and N-Q, respectively. White arrowheads indicate cells co-expressing all three genes. The region enclosed by dashed boxes in panels I-M is enlarged in panels N-R. Panels I-M present maximum-intensity projections of confocal z-stacks, whereas panels N-R show single optical sections. Cell counts and co-expression analyses (S–T) were derived from these FISH datasets by nuclear segmentation to define individual cells and assignment of transcript spot localizations to segmented nuclei (See Supplementary Fig. S6 for an example workflow/output). (S) Co-expression pattern of *PT1*, *PT2*, and *PP1*. Numbers in the diagram indicate numbers of cells expressing each gene combination. A total of 978 cells (including triple-negative cells) in the pineal gland were analyzed. The Venn diagram summarizes only cells positive for at least one marker (n = 471). Pairwise co-expression between *PT1*, *PT2*, and *PP1* was assessed using Fisher’s exact test (two-sided) on 2×2 contingency tables of cell-level positivity/negativity. To control the familywise error rate across the three pairwise tests, p-values were adjusted using the Holm procedure. Effect sizes are reported as odds ratios (OR) with 95% Wald confidence intervals (CI). *PT1* vs *PT2*: OR = 29.16 (95% CI 18.92–44.93), p = 3.59×10⁻^78^, Holm-adjusted p = 1.08×10⁻^77^ *PP1* vs *PT2*: OR = 12.41 (95% CI 9.031–17.05), p = 2.20×10⁻^64^, Holm-adjusted p = 4.40×10⁻^64^. *PP1* vs *PT1*: OR = 13.25 (95% CI 9.011–19.49), p = 3.49×10⁻^51^, Holm-adjusted p = 3.49×10⁻^51^. (T) Transcript numbers of *PT1*, *PT2*, and *PP1* per cell. For analysis of FISH images, cells with two or more fluorescent spots were classified as expressing the respective gene. Statistical analysis was performed using the Kruskal-Wallis test, followed by the Dunn-Holland-Wolfe test for post-hoc pairwise comparisons. (U) Quantification of expression levels of *PT1*, *PT2*, and *PP1* by RT-qPCR. Statistical analysis was performed using one-way ANOVA test, followed by the Tukey-Kramer’s test for post-hoc pairwise comparisons. Scale bars: 100 μm (A, I); 5 μm (N).

## Discussion

Here, we showed that (1) TGD paralogs of PT (PT1 and PT2) are retained in a few species including the red piranha, and while most teleosts have only PT1, the Mexican tetra and catfishes have only PT2, (2) spectral properties are comparable between TGD paralogs of PT (PT1 and PT2), (3) in a heterologous cell-based assay, red piranha PT1 increases cellular cAMP in response to light, whereas PT2 decreases it, and (4) in the pineal gland of the red piranha, *PT1*, *PT2*, and *PP1* are co-expressed. Previous studies have primarily focused on differences in spectral properties and/or expression patterns of TGD paralogous opsin pairs [5–7]. This study illustrates the importance of surveying differences in intracellular signaling properties to understand how TGD contributed to functional diversification of opsins.

### Evolutionary scenario explaining how PTs may have diversified after TGD

Here, we show that TGD paralogs of PT (PT1 and PT2) are retained in a few characins, including the red piranha. While most teleosts have only PT1, the Mexican tetra, which is a characin, and catfishes have only PT2. Judging from phylogenetic positions of characins and catfishes [19,25], PT2 must have been lost several times independently in evolution (Supplementary Fig. S4). Furthermore, considering that they are phylogenetically close [19,25], a reversal of this "PT2 preferentially lost tendency" may have occurred in this lineage (Supplementary Fig. S4). How this can be explained is an intriguing question. One possibility is that diversification of PTs was gradual. PT1 and PT2 may have remained functionally redundant long after TGD and may have diversified relatively recently. Indeed, our recent studies of RH1 in ray-finned fishes suggest that diversification processes can be gradual after duplication of opsins. RH1 experienced retrotransposition/retroduplication in the early phase of ray-finned fish evolution, resulting in one paralog with introns and the other without [26] (Fig. 1A). Both are expressed mainly in the retina in non-teleost fishes with similar absorption maxima and decay rates of meta II intermediate [26]. In teleosts, while intronless RH1s are expressed in the retina, RH1s with introns (exo-rhodopsin) are expressed in the pineal gland, suggesting acquisition of this novel expression pattern in early teleost evolution [27,28]. Exo-rhodopsin proteins have a shorter lifetime of meta II compared to those of intronless paralogs, presumably reflecting acquisition of novel functions optimized for pineal photoreception under continuous bright light conditions [12,28,29].

Recently, it was suggested that functionally redundant paralogs can be retained in a vertebrate genome for extended evolutionary periods [30]. This may support our gradual diversification scenario.

The reason why PT2s are preferentially lost in most teleosts is not necessarily clear in this scenario. Biased retention/loss between TGD paralogs reported in [19] may provide one possible explanation.

To further understand opsin evolution, it will be desirable to continue similar cell culture-based assays for other species PTs, especially non-teleost PTs. Such studies may provide insights into evolution of vertebrate photoreception, considering that (1) PT comprises one of the subfamilies of vertebrate visual and non-visual opsin, which is important in understanding acquisition of sophisticated photoreception systems unique to vertebrates [1,2], (2) the response to light of PT in cultured cells has not been reported in other species [13,21,22], and (3) although a jellyfish opsin can increase cellular cAMP in response to light [31], to our knowledge, opsins that can only increase cellular cAMP in response to light have never been reported in bilaterians.

This study illustrates the importance of characterizing opsins in teleosts other than classical model organisms such as the zebrafish and medaka. Although the red piranha and the zebrafish are phylogenetically close [19,25], several differences in their opsin repertoires have been reported [6,32]. Furthermore, we detected dispersed mRNA expression of *PT1* in the optic tract of the red piranha (Supplementary Fig. S5). To our knowledge, *PT* expression outside the pineal gland or related organs has rarely been reported. Whether this expression is evolutionarily conserved and whether it is involved in molecular properties characteristic of PT is an intriguing topic. Considering these, the red piranha will serve as a valuable experimental system in future opsin studies.

### Co-expression of PT & PP and its possible function in the pineal gland

We found that *PT1* and *PT2* are co-expressed with *PP1* in the red piranha pineal gland. Pineal co-expression of UV-sensitive PP and green-sensitive PT is reported in zebrafish and the green iguana, suggesting an ancient origin [21,24]. Why they are co-expressed is not clear at present. One possibility is that they are involved in pineal chromatic opponency, although a zebrafish study suggests that PP1, which is UV-sensitive in its 11-*cis* retinal bound form and visible light-sensitive in its all-*trans* retinal bound form, is likely responsible for chromatic opponency alone [21]. Another, non-mutually exclusive possibility is that UV-sensitive PP-positive cells are multifunctional, and that PT has some unknown functions there. Our recent extensive survey supports this idea. That is, retention/loss of UV-sensitive PP1, but not visual-light-sensitive PP2, seems tightly linked to that of PT. Of 112 teleost species we surveyed, (1) 89 retain PP1, PP2, and PT, (2) nine species have lost all three, presumably reflecting reduced dependency on pineal photoreception, (3) nine species, including the Siamese fighting fish (*Betta splendens*), have lost PP2, but retain PP1 and PT, (4) four species, including medaka, have lost both PP1 and PT, but retain PP2, and (5) uncoupled retention/loss of PP1 and PT was observed only in the European eel (*A. anguilla*), which retains PP1, but lost PT [17]. Although the photophysiological role of PT is not very clear at present, its most prominent molecular property is the longest-wavelength sensitivity among vertebrate non-visual opsins reported [13,21,22]. Considering these characteristics, determining the nature of possible relationships between green-sensitive PT and UV-sensitive PP in the same cells will be important in order to understand evolutionarily conserved functions of the pineal gland. Finally, how co-expressed PT1 and PT2, which exhibit opposite light responses, at least in cultured cells, function together in pineal cells will be an interesting subject for future study. Together with the generally higher expression of *PT2* and lower expression of *PT1*, as well as heterogeneous expression profiles, their co-expression may enable fine-tuning of cellular responses across a wide range of light intensities. Their expression levels may also be regulated by factors such as environmental cues and developmental stages.

## Conclusions

Here, we showed that (1) TGD paralogs of PT (PT1 and PT2) are retained in a few species, including the red piranha, and that while most teleosts have only PT1, the Mexican tetra and catfishes have only PT2, (2) spectral properties are comparable between PT1s and PT2s, (3) in a heterologous cell-based assay, red piranha PT1 increases cellular cAMP in response to light, whereas PT2 decreases it, while neither induced a clear Ca²⁺ response under the conditions tested, and (4) in the pineal gland of the red piranha, *PT1*, *PT2*, and *PP1* are co-expressed. This study provides valuable insights into how TGD contributed to functional diversification of opsins.

## Methods

### Molecular phylogenetic analyses

Amino acid sequences were aligned using MAFFT v7.525 [33] with the option ’-linsi.’ Gaps and unaligned regions were removed using gblocks [34]. Molecular phylogenetic analyses were then performed using Bayesian Inference (BI) and the Maximum-Likelihood (ML) method with the LG+G model. BI analyses were performed using MrBayes v3.2.3 [35] with the following parameters: ngen=200000000 printfreq=10000 samplefreq=100 nchains=4 temp=0.2 checkfreq=50000 diagnfreq=500000 stopval=0.01 stoprule=yes. The first 25% of these trees were discarded as ‘burn-in.’ Convergence of each run was assessed by plotting the log-likelihood. ML analyses were performed using RAxML-NG v1.0.2 [36] with 1000 bootstrap pseudoreplications.

### Preparation of recombinant proteins and spectroscopic measurements

cDNAs encoding red piranha (*Pygocentrus nattereri*) PT1 (XP_017545252) and PT2 (XP_017567667), Mexican tetra (*Astyanax mexicanus*) PT2 (XP_007252482), and Japanese catfish (*Silurus asotus*) PT2 (KAI5617949) were tagged with the epitope sequence of the anti-bovine rhodopsin monoclonal antibody, Rho1D4, at their C-termini and were introduced into the mammalian expression vector, pCAGGS. Plasmids were transfected into HEK293S cells using the calcium-phosphate method. Twenty-four hours after transfection, 11-*cis* retinal was added to the culture medium to a final concentration of 5 µM. Cells were then kept in the dark and harvested 48 h post-transfection. Reconstituted parietopsins were solubilized in 1% n-dodecyl-β-D-maltoside (DDM) prepared in buffer A (50 mM HEPES, 140 mM NaCl, pH 6.5). UV-visible absorption spectra of solubilized samples in the presence of 100 mM NH_2_OH were recorded at 0 ± 0.1°C with a UV-2600 spectrophotometer (Shimadzu) using a quartz cuvette (2 mm width, 1 cm path length). Samples were irradiated through an R-62 cutoff filter from a 1-kW tungsten–halogen lamp (Master HILUX-HR, Rikagaku Seiki).

### Animals

Red piranhas (*P. nattereri*, body length: ∼ 5 cm) were purchased from local pet shops.

### Isolation of opsin cDNAs

Total RNA from the red piranha pineal gland was extracted and purified using a Direct-zol RNA kit (Zymo Research), according to the manufacturer’s instructions.

First-strand cDNA was synthesized using a PrimeScript IV 1st strand cDNA Synthesis Mix (Takara). Fragments of red piranha parietopsin and parapinopsin genes (*PT1*, *PT2*, and *PP1*) were amplified by PCR from first-strand cDNA using the following primer pairs: *PT1*, PN_PT1_Fw (5’-TAAACACACCAACAAACACAGCGCC-3’) and PN_PT1_Rv (5’-TCATCCGTAAGGAGCCCATCTCCAT-3’); *PT2*, PN_PT2_Fw (5’-ACATGTGGAACGAAGCACAGAGGAG-3’) and PN_PT2_Rv (5’-GGAGACAATAGCATAGGGCAGCCAG-3’) ; *PP1*, PN_PP1_Fw (5’-CAAGGATAGGCTACACCGTCTTGGC-3’) and PN_PP1_Rv (5’-TTCCGCAGAACAGATATGGCAGAGC -3’).

### Preparation of tissue sections

Red piranhas were euthanized by immersion in 0.1% 2-phenoxyethanol, followed immediately by decapitation. To improve reagent penetration, portions of the cranial bone were removed, and heads were fixed overnight at 4°C in phosphate-buffered saline (PBS) containing 4% paraformaldehyde. Specimens were then decalcified for five days at 4°C in 10% or 14% EDTA (pH 8.0). After cryoprotection in PBS with 20% sucrose overnight at 4°C, tissues were embedded in OCT compound (Tissue-Tek) and snap-frozen at -80°C. Frozen tissues were sectioned at 15 µm, mounted on glass slides, air-dried, and stored at -20°C until use.

### Cellular cAMP response assay using GloSensor

The coding sequence of human codon–optimized GloSensor-22F cAMP biosensor (Glo22F) was synthesized by Thermo Scientific and inserted into the mammalian expression vector, pCAGGS [37]. HEK293T cells were seeded into white-walled 96-well plates (MS-8096W, Sumitomo Bakelite) at 20,000 cells per well in Dulbecco’s modified Eagle’s medium/F-12 (Wako) supplemented with 10% FBS and cultured at 37°C in a CO_2_ incubator. After 24 hours, cells were transfected with expression plasmids encoding the opsin and Glo22F using polyethyleneimine (PEI MAX, Polyscience). A mixture of 10 μL Opti-MEM (ThermoFisher Scientific), 0.4 μg PEI MAX, and 0.1 μg total of plasmids was applied to each well of a 96-well plate. The ratio of plasmids of opsin to Glo22F was 1:1. Six to eight hours after transfection, medium was replaced with fresh medium containing 2 μM 11-*cis* retinal. On the following day, medium was replaced with phenol red–free L-15 medium (ThermoFisher Scientific) containing 10% FBS and 2 mM D-luciferin (Goryo Chemical, Sapporo, Japan). Incubation and luminescence measurements were performed at 25 ± 1°C in an air-conditioned room. After 2 h of incubation in the dark, luminescence was measured using a microplate luminometer (Veritas, Turner Biosystems). To elevate intracellular cAMP levels, forskolin was added at a final concentration of 2 μM. Cells were stimulated with a homemade green LED array (8 μW mm^−2^; peak irradiance at 500 nm) for 30 seconds.

### Aequorin luminescence–based calcium assay

A luminescent calcium assay using aequorin as an indicator was performed according to previous methods [38,39]. HEK293T cells were seeded into white-walled 96-well plates (SPL Life sciences) at a density of 60,000 cells per well in Dulbecco’s modified Eagle’s medium/F-12 (Thermo Fisher Scientific) containing 10% FBS. After incubation for 24 hr, cells were transfected with 50 ng of opsin plasmid and 50 ng of aequorin plasmid per well using polyethylenimine (PEI MAX, Polysciences). After 6 hr of incubation, medium was replaced with fresh medium containing 2.5 μM 11-*cis* retinal. After overnight incubation, medium was replaced with CO_2_-independent medium (Thermo Fisher Scientific) containing 10% FBS and 10 μM coelenterazine h (Fuji Film). After 2 hr of incubation in the dark, luminescence was measured using a multimode microplate reader (GloMax Explorer, Promega). Cells were stimulated with a fluorescence excitation system attached to the equipment (peak irradiance at 520 nm).

### Chromogenic *in situ* hybridization

*In situ* hybridization was performed as in a previous study [28]. Digoxigenin-labeled sense and antisense riboprobes were synthesized from cDNAs flanked by T7 and T3 promoter sequences, which were inserted into pBluescript II KS+. Unless otherwise specified, all subsequent steps were carried out at room temperature. Tissue sections were sequentially immersed in 4% PFA in PBS for 15 min, 100% methanol for 30 min, PBS for 5 min, Proteinase K (0.5 μg/mL) in Tris–EDTA buffer (50 mM Tris–HCl, 5 mM EDTA, pH 7.6) for 15 min, 4% PFA in PBS for 15 min, dimethyl dicarbonate-treated water for 30 s, acetylation buffer (0.27% (v/v) acetic anhydride, 100 mM triethanolamine, pH 8.0) for at least 30 min, PBS for 5 min, and finally hybridization buffer (750 mM NaCl, 75 mM sodium citrate, 0.4 mg/mL yeast RNA, 0.1 mg/mL heparin sodium, 1 × Denhardt’s solution, 0.1% (v/v) Tween, 0.1% (w/v) CHAPS, 5 mM EDTA, 70% (v/v) formamide) at 65°C for 2 ∼ 3 h. Digoxigenin-labelled riboprobes (final concentration: 0.17 μg/mL) diluted in hybridization buffer were then applied to tissue sections and incubated at 65°C for approximately 40 h. After hybridization, tissue sections were washed in 1 × SSC buffer (150 mM NaCl, 15 mM sodium citrate, pH 7.0) containing 50% formamide at 65°C for 15 min and 1 h, followed by washing in 0.2 × SSC buffer at 65°C for 1 h, and then in MABT (100 mM maleate, 150 mM NaCl, 0.1% Tween 20, pH 7.5) three times for 30 min each. After washing, tissue sections were blocked with PBS containing 1% (w/v) bovine serum albumin, 10% (v/v) sheep normal serum, and 0.08% (v/v) Triton-X100 for 30 min and then incubated with alkaline phosphatase-conjugated anti-digoxigenin Fab fragment (1:2000 dilution; Roche) overnight at 4°C. Tissue sections were then washed three times for 30 min each with MABT and twice for 5 min each with AP reaction buffer (100 mM Tris–HCl, 50 mM MgCl2, 100 mM NaCl, and 0.1% Tween 20, pH 9.5). Color development was performed with 50 μg/mL nitro blue tetrazolium, 175 μg/mL 5-bromo-4-chloro-3-indolyl phosphate, 1 mM levamisole hydrochloride, and 5% (w/v) polyvinyl alcohol (M.W. 85,000∼124,000, Sigma-Aldrich) in AP reaction buffer at 28°C for 2 days. Finally, sections were immersed for 5 min in PBS containing 0.1% Triton X 100, rinsed with water, counterstained with Nuclear Fast Red, washed in tap water for 5 min, dehydrated twice in isopropanol for 2 min, and coverslipped with VectaMount Express mounting medium (Vector Laboratories).

### Fluorescence *in situ* hybridization

Simultaneous detection of *PT1*, *PT2*, and *PP1* mRNAs in red piranha pineal gland was performed with signal amplification by exchange reaction–fluorescence *in situ* hybridization (SABER-FISH) [40]. Probe sequences were designed using the blockParse.py script from the OligoMiner package on the coding sequences of *PT1*, *PT2*, and *PP1* [41]. Primer exchange reactions were performed following previous reports. Fluorescent signals were amplified and detected with two rounds of branching as follows: tissue sections were treated with 25 μg/mL pepsin in 0.01 M HCl at 37°C for 20 min, followed by three rinses in PBS containing 0.1% Tween20 (PBST). They were pre-hybridized in Prehyb (40% formamide, 2×SSC, 1% Tween20, 1×Denhardt’s solution, 100 μg/mL heparin sodium, and 0.5 μM random 20 mer oligo DNA) at 43°C for 10 min, and then they were incubated with respective probes at 1 μg/mL in Hyb1.1 (Prehyb containing 10% dextran sulfate, FUJIFILM Wako Pure Chemical Corporation 197-09984) at 43°C for 16 h. Post-hybridization washes were performed at 43°C in Whyb (40% formamide, 2×SSC, 1% Tween20) for two 30-min washes, followed by 2×SSCT. For signal amplification, two rounds were performed at 37°C: hybridization with extended branch DNA (1 μg/mL in Hyb1.1, 12 h), two 30-min washes in Whyb, and a rinse in 2× SSCT. The branch-oligo sequence was exchanged between rounds. Fluorophore-conjugated oligos (0.1 μM) diluted in Hyb2.1 (0.1% Tween20, 1× Denhardt’s solution, 100 μg/mL heparin sodium, and 0.5 μM random 20-mer DNA oligonucleotide in PBS) were added for 1 h at 37°C. Finally, sections were counterstained with DAPI (0.2 µg/mL), rinsed in PBST, and coverslipped with homemade polyvinyl alcohol/glycerol mounting medium. Oligonucleotide sequences used in SABER-FISH are listed in Supplementary Table S2. Confocal fluorescence images were collected with a Carl Zeiss LSM 780 laser scanning confocal microscope system at the Central Research Laboratory, Okayama University Medical School.

### Quantitative analysis of FISH images

Z-stack FISH images acquired by confocal microscopy were quantitatively analyzed as follows. First, nuclei were segmented from the DAPI channel with cellpose-SAM [42]. Next, fluorescent spots corresponding to *PT1*, *PT2*, and *PP1* transcripts were detected in their respective channels using RS-FISH, with parameters set to sigma=1.5 and threshold=0.007 [43]. Detected spots were assigned to the nearest segmented nucleus based on 3D (x–y–z) distances. An overview of the analysis workflow and representative outputs are shown in Supplementary Fig. S6. The spot-localization table including x–y–z coordinates and the per-nucleus summary table reporting the number of assigned spots per gene are available in Supplementary Data 2. To filter out potential noise from single spots, only cells containing two or more spots for a specific gene were classified as expressing. All statistical analyses were conducted using Igor Pro 9.05 (64-bit) and 10.00 (64-bit).

### Reverse transcription-quantitative PCR (RT-qPCR)

Total RNA from red piranha pineal gland was extracted and purified using a FastGene RNA Premium Kit (Nippon Genetics) according to the manufacturer’s instructions. To generate standard curves that incorporate reverse-transcription efficiency, synthetic RNA standards for *PT1*, *PT2*, and *PP1* were prepared as follows. Gene fragments of *PT1*, *PT2*, and *PP1* inserted into the EcoRV site of pBluescript KS+ were amplified using T7 and T3 primers, and PCR products were purified. Using purified PCR products as templates, RNA was synthesized with Thermo T7 RNA Polymerase (TRL-201; TOYOBO). The transcription reaction contained 2 µg template DNA, 25 µL 10× transcription buffer (TOYOBO), 10 mM each NTP mix (GeneACT), 2.5 µL of 1 M DTT, 2.5 µL pyrophosphatase (Takara), 1 µL of 2 M MgCl₂, and 3.125 µL T7 polymerase, and was incubated at 37°C for 4 h. After ethanol precipitation, RNA concentration was measured, and a serial dilution series corresponding to 10 –10² molecules/μL was prepared and used as the template for reverse transcription. First-strand cDNA was synthesized using a PrimeScript FAST RT reagent Kit with gDNA Eraser (Takara) according to the manufacturer’s instructions. qPCR was performed on a LightCycler Nano (Roche Diagnostics) using THUNDERBIRD Next SYBR qPCR Mix (Toyobo; QPX-201) according to the manufacturer’s instructions. qPCR primer sequences and amplicon sizes are listed in Supplementary Table S3. For each gene, Cq values were normalized across samples using the geometric mean of the reference genes (*ef1a*, *rpl7*, and *actb2*) to correct for between-sample variation, and transcript molecule numbers in the original samples were then estimated using the standard curves generated from the synthetic RNA standards (Supplementary Fig. S7). Statistical analysis was performed using Igor Pro 10.00 (64-bit).

## Abbreviations

TGD: Teleost whole-genome duplication
PP: Parapinopsin
VA: Vertebrate Ancient
LWS: Long wavelength-sensitive opsin
PT: Parietopsin
GPCR: G-protein-coupled receptor
BI: Bayesian Inference
ML: Maximum-Likelihood
DDM: n-dodecyl-β-D-maltoside
PBS: Phosphate-buffered saline
Glo22F: GloSensor-22F cAMP biosensor
SABER: Signal amplification by exchange reaction
FISH: Fluorescence *in situ* hybridization
PBST: PBS containing 0.1% Tween20
SWS: short wavelength-sensitive opsin
RH2: rhodopsin-like 2

## Declarations

### Ethics approval and consent to participate

Use of animals in these experiments conformed to guidelines established by the Japan Ministry of Education, Culture, Sports, Science and Technology. The experimental design was approved by the Animal Care and Use Committee, Okayama University.

### Consent for publication

Not applicable.

### Availability of data and materials

Data supporting this article are available in the article and in online supplementary material.

### Competing interests

The authors declare no competing interests.

### Funding

This work was supported in part by Japan Society for the Promotion of Science (JSPS) Grants-in-Aid for Scientific Research to KSato (23K05850), TY (24K09530), and TGK (21K19280, 23K27185, 23H02492), Japan Agency for Medical Research and Development (AMED) CREST to TY (22gm1510007), Frontier Photonic Sciences Project of National Institutes of Natural Sciences to KSato (01212503), grants from the Japan Health Foundation and the Japan Foundation for Applied Enzymology to KSato, a grant from the Naito Foundation to TY.

### Authors’ contributions

KSato, TY, TGK, and FG designed the research. KSato, TY, and KSakai performed the experiments. FG performed the molecular evolutionary studies. KSato, TY, KSakai, HO, TGK, and FG analyzed the data. KSato, TY, and FG wrote the manuscript with editing by all authors.

## Acknowledgements

We thank Prof. Robert S. Molday for the generous gift of a Rho1D4-producing hybridoma, Prof. S. Koike for HEK293T cell line, and Prof. Jeremy Nathans for providing the HEK293S cell line. The manuscript was edited by Dr. Steven D. Aird (https://www.sda-technical-editor.org).

## Notes

### Competing Interest Statement

The authors have declared no competing interest.

### Summary of Updates

New experimental data added; main text revised; Figures revised; authors updated; Supplemental files updated.

## References

1. Terakita A. The opsins. Genome Biol. 2005;6:213.

2. Shichida Y, Matsuyama T. Evolution of opsins and phototransduction. Philos Trans R Soc Lond B Biol Sci. 2009;364:2881–95.

3. Ramirez M, Pairett A, Pankey M, Serb J, Speiser D, Swafford A, et al. The last common ancestor of most bilaterian animals possessed at least 9 opsins. Genome Biol Evol. 2016;8:3640–52.

4. Beaudry FEG, Iwanicki TW, Mariluz BRZ, Darnet S, Brinkmann H, Schneider P, et al. The non-visual opsins: eighteen in the ancestor of vertebrates, astonishing increase in ray-finned fish, and loss in amniotes. J Exp Zool B Mol Dev Evol. 2017;328:685–96.

5. Koyanagi M, Wada S, Kawano-Yamashita E, Hara Y, Kuraku S, Kosaka S, et al. Diversification of non-visual photopigment parapinopsin in spectral sensitivity for diverse pineal functions. BMC Biol. 2015;13:73.

6. Liu D-W, Wang F-Y, Lin J-J, Thompson A, Lu Y, Vo D, et al. The cone opsin repertoire of osteoglossomorph fishes: Gene loss in mormyrid electric fish and a long wavelength-sensitive cone opsin that survived 3R. Mol Biol Evol. 2019;36:447–57.

7. Kojima D, Torii M, Fukada Y, Dowling JE. Differential expression of duplicated VAL-opsin genes in the developing zebrafish. J Neurochem. 2008;104:1364–71.

8. Davies WIL, Zheng L, Hughes S, Tamai TK, Turton M, Halford S, et al. Functional diversity of melanopsins and their global expression in the teleost retina. Cell Mol Life Sci. 2011;68:4115–32.

9. Nakamura Y, Yasuike M, Mekuchi M, Iwasaki Y, Ojima N, Fujiwara A, et al. Rhodopsin gene copies in Japanese eel originated in a teleost-specific genome duplication. Zoological Lett. 2017;3:18.

10. Yamaguchi K, Koyanagi M, Kuraku S. Visual and nonvisual opsin genes of sharks and other nonosteichthyan vertebrates: Genomic exploration of underwater photoreception. J Evol Biol. 2021;34:968–76.

11. Fujiyabu C, Sato K, Ohuchi H, Yamashita T. Diversification processes of teleost intron-less opsin genes. J Biol Chem. 2023;299:104899.

12. Morrow JM, Lazic S, Dixon Fox M, Kuo C, Schott RK, de A Gutierrez E, et al. A second visual rhodopsin gene, rh1-2, is expressed in zebrafish photoreceptors and found in other ray-finned fishes. J Exp Biol. 2017;220(Pt 2):294–303.

13. Su C-Y, Luo D-G, Terakita A, Shichida Y, Liao H-W, Kazmi MA, et al. Parietal-eye phototransduction components and their potential evolutionary implications. Science. 2006;311:1617–21.

14. FoàA, Basaglia F, Beltrami G, Carnacina M, Moretto E, Bertolucci C. Orientation of lizards in a Morris water-maze: roles of the sun compass and the parietal eye. J Exp Biol. 2009;212:2918–24.

15. Nishimura T, Okano H, Tada H, Nishimura E, Sugimoto K, Mohri K, et al. Lizards respond to an extremely low-frequency electromagnetic field. J Exp Biol. 2010;213:1985–90.

16. Solessio E, Engbretson GA. Antagonistic chromatic mechanisms in photoreceptors of the parietal eye of lizards. Nature. 1993;364:442–5.

17. Gyoja F, Sato K, Yamashita T, Kusakabe TG. An extensive survey of vertebrate-specific, nonvisual opsins identifies a novel subfamily, Q113-bistable opsin. Genome Biol Evol. 2025;17:evaf032.

18. Parey E, Louis A, Cabau C, Guiguen Y, Roest Crollius H, Berthelot C. Synteny-guided resolution of gene trees clarifies the functional impact of whole-genome duplications. Mol Biol Evol. 2020;37:3324–37.

19. Parey E, Louis A, Montfort J, Guiguen Y, Crollius HR, Berthelot C. An atlas of fish genome evolution reveals delayed rediploidization following the teleost whole-genome duplication. Genome Res. 2022;32:1685–97.

20. Nakatani Y, McLysaght A. Genomes as documents of evolutionary history: a probabilistic macrosynteny model for the reconstruction of ancestral genomes. Bioinformatics. 2017;33:i369–78.

21. Wada S, Shen B, Kawano-Yamashita E, Nagata T, Hibi M, Tamotsu S, et al. Color opponency with a single kind of bistable opsin in the zebrafish pineal organ. Proc Natl Acad Sci U S A. 2018;115:11310–5.

22. Sakai K, Imamoto Y, Su C-Y, Tsukamoto H, Yamashita T, Terakita A, et al. Photochemical nature of parietopsin. Biochemistry. 2012;51:1933–41.

23. Wada S, Kawano-Yamashita E, Sugihara T, Tamotsu S, Koyanagi M, Terakita A. Insights into the evolutionary origin of the pineal color discrimination mechanism from the river lamprey. BMC Biol. 2021;19:188.

24. Wada S, Kawano-Yamashita E, Koyanagi M, Terakita A. Expression of UV-sensitive parapinopsin in the iguana parietal eyes and its implication in UV-sensitivity in vertebrate pineal-related organs. PLoS One. 2012;7:e39003.

25. Betancur-R R, Wiley EO, Arratia G, Acero A, Bailly N, Miya M, et al. Phylogenetic classification of bony fishes. BMC Evol Biol. 2017;17:162.

26. Fujiyabu C, Sato K, Utari NML, Ohuchi H, Shichida Y, Yamashita T. Evolutionary history of teleost intron-containing and intron-less rhodopsin genes. Sci Rep. 2019;9:10653.

27. Mano H, Kojima D, Fukada Y. Exo-rhodopsin: a novel rhodopsin expressed in the zebrafish pineal gland. Brain Res Mol Brain Res. 1999;73:110–8.

28. Fujiyabu C, Gyoja F, Sato K, Kawano-Yamashita E, Ohuchi H, Kusakabe TG, et al. Functional diversification process of opsin genes for teleost visual and pineal photoreceptions. Cell Mol Life Sci. 2024;81:428.

29. Tarttelin EE, Fransen MP, Edwards PC, Hankins MW, Schertler GFX, Vogel R, et al. Adaptation of pineal expressed teleost exo-rod opsin to non-image forming photoreception through enhanced Meta II decay. Cell Mol Life Sci. 2011;68:3713–23.

30. Fujimori C, Sugimoto K, Ishida M, Yang C, Kayo D, Tomihara S, et al. Long-lasting redundant *gnrh1/3* expression in GnRH neurons enabled apparent switching of paralog usage during evolution. iScience. 2024;27:109304.

31. Koyanagi M, Takano K, Tsukamoto H, Ohtsu K, Tokunaga F, Terakita A. Jellyfish vision starts with cAMP signaling mediated by opsin-G(s) cascade. Proc Natl Acad Sci U S A. 2008;105:15576–80.

32. Musilova Z, Cortesi F. The evolution of the green-light-sensitive visual opsin genes (RH2) in teleost fishes. Vision Res. 2023;206:108204.

33. Katoh K, Misawa K, Kuma K-I, Miyata T. MAFFT: a novel method for rapid multiple sequence alignment based on fast Fourier transform. Nucleic Acids Res. 2002;30:3059–66.

34. Talavera G, Castresana J. Improvement of phylogenies after removing divergent and ambiguously aligned blocks from protein sequence alignments. Syst Biol. 2007;56:564–77.

35. Huelsenbeck JP, Ronquist F. MRBAYES: Bayesian inference of phylogenetic trees. Bioinformatics. 2001;17:754–5.

36. Kozlov AM, Darriba D, Flouri T, Morel B, Stamatakis A. RAxML-NG: a fast, scalable and user-friendly tool for maximum likelihood phylogenetic inference. Bioinformatics. 2019;35:4453–5.

37. Inoue A, Raimondi F, Kadji FMN, Singh G, Kishi T, Uwamizu A, et al. Illuminating G-Protein-Coupling Selectivity of GPCRs. Cell. 2019;177:1933–47.e25.

38. Bailes HJ, Lucas RJ. Human melanopsin forms a pigment maximally sensitive to blue light (λmax ≈ 479 nm) supporting activation of G(q/11) and G(i/o) signalling cascades. Proc Biol Sci. 2013;280:20122987.

39. Sakai K, Ikeuchi H, Fujiyabu C, Imamoto Y, Yamashita T. Convergent evolutionary counterion displacement of bilaterian opsins in ciliary cells. Cell Mol Life Sci. 2022;79:493.

40. Kishi JY, Lapan SW, Beliveau BJ, West ER, Zhu A, Sasaki HM, et al. SABER amplifies FISH: enhanced multiplexed imaging of RNA and DNA in cells and tissues. Nat Methods. 2019;16:533–44.

41. Beliveau BJ, Kishi JY, Nir G, Sasaki HM, Saka SK, Nguyen SC, et al. OligoMiner provides a rapid, flexible environment for the design of genome-scale oligonucleotide in situ hybridization probes. Proc Natl Acad Sci U S A. 2018;115:E2183–92.

42. Pachitariu M, Rariden M, Stringer C. Cellpose-SAM: superhuman generalization for cellular segmentation. bioRxiv. 2025. 10.1101/2025.04.28.651001

43. Bahry E, Breimann L, Zouinkhi M, Epstein L, Kolyvanov K, Mamrak N, et al. RS-FISH: precise, interactive, fast, and scalable FISH spot detection. Nat Methods. 2022;19:1563–7.

44. Musilova Z, Salzburger W, Cortesi F. The Visual Opsin Gene Repertoires of Teleost Fishes: Evolution, Ecology, and Function. Annu Rev Cell Dev Biol. 2021;37:441–68.

45. Parey E, Louis A, Montfort J, Bouchez O, Roques C, Iampietro C, et al. Genome structures resolve the early diversification of teleost fishes. Science. 2023;379:572–5.

