## Supplementary Figures for "How opsins diversified after the teleost whole-genome duplication: Insights from two parietopsins of the red piranha, *Pygocentrus nattereri*"

**Supplementary Figure S1**

**Comparison of spectral property of PT1 and PT2.**

Spectra shown in Fig. 3 were normalized to be ~ 1.0 at the positive maximum. Normalized spectra of red piranha PT1 (black curve), red piranha PT2 (red curve), Mexican tetra PT2 (green curve) and Japanese catfish PT2 (blue curve) are compared.

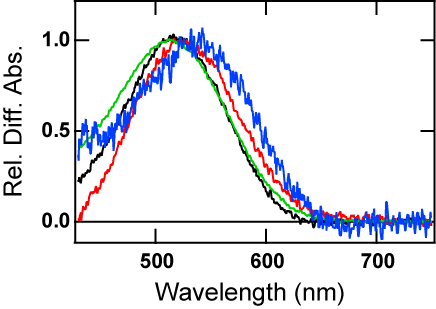

**Supplementary Figure S2**

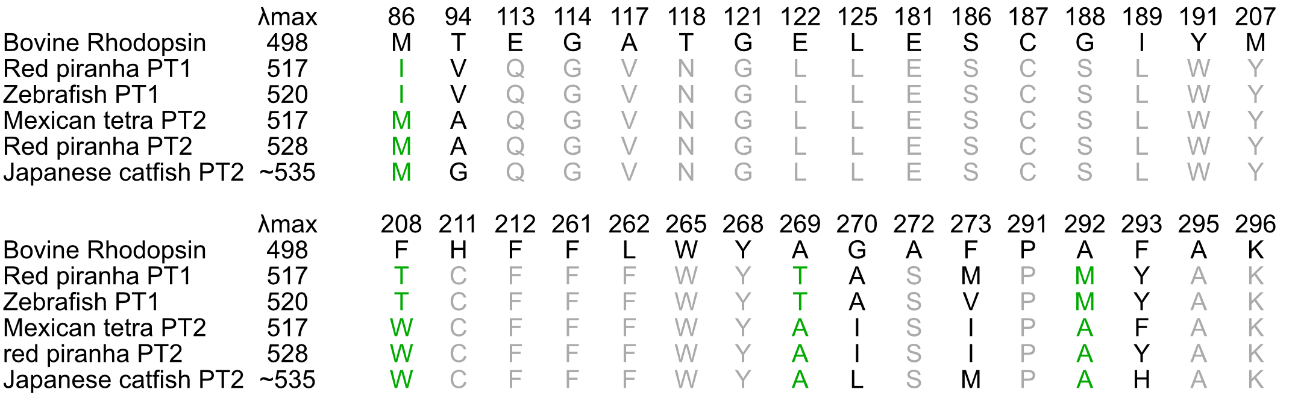

**Comparison of amino acid residues surrounding the retinal chromophore in parietopsins.**

Amino acid residues located within 6 Å of 11-*cis*-retinal were compared among bovine rhodopsin, red piranha PT1, zebrafish PT1, Mexican tetra PT2, Japanese catfish PT2, and red piranha PT2. Structural models of parietopsins were generated using AlphaFold3 and aligned to bovine rhodopsin (PDB: 1U19). Residue positions are shown according to bovine rhodopsin numbering. Green letters indicate residues that appear to be conserved within either the PT1 or PT2 lineage in this limited comparison, whereas gray letters indicate residues conserved between PT1 and PT2. Although several lineage-associated residues were observed, no single residue or simple motif clearly correlated with the differences in absorption maxima among the examined parietopsins.

**Supplementary Figure S3**

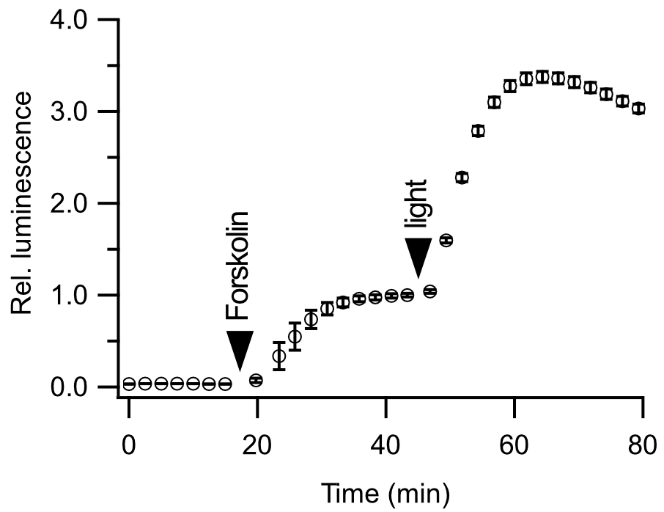

**Light-dependent cAMP response of red piranha PT1 after forskolin stimulation.**
Time course of relative luminescence in the GloSensor 22F cAMP assay in cells expressing red piranha PT1. Forskolin was added during luminescence recording at a final concentration of 2 μM, as indicated, and cells were subsequently stimulated with light. Light stimulation induced a further increase in luminescence in PT1-expressing cells after forskolin-mediated elevation of intracellular cAMP. The light-induced increase in luminescence in PT1-expressing cells was larger in the presence of forskolin than in its absence. The mechanism underlying this difference remains unclear, but it may be explained by the dynamic range of cAMP probe GloSensor 22F under the assay conditions. Data are shown as mean ± SEM (n = 3). Relative luminescence was normalized to the baseline before light stimulation.

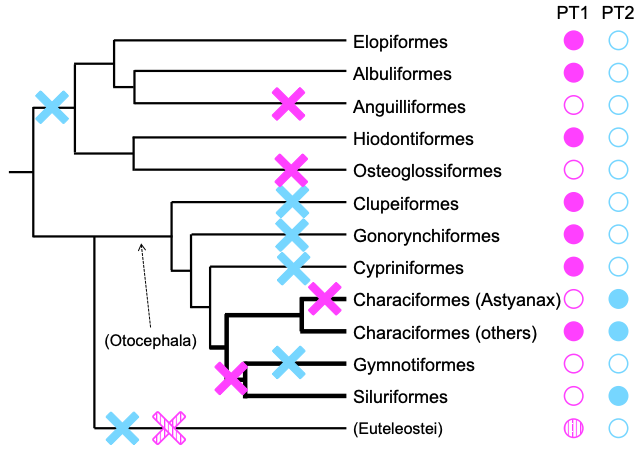
**Supplementary Figure S4**

**A reversal of the "PT2 preferentially lost tendency" may have occurred during teleost evolution.**

Most teleosts have lost PT2 while retaining PT1. However, PT1 may tend to be lost more than PT2 in the clade containing Characiformes (characins) and Siluriformes (catfishes). Deduced loss events of PT1 (magenta) and PT2 (pale blue) are shown on a phylogenetic tree, based on Gyoja et al. (2025) and Figure 1B. Filled Xs indicate that all species surveyed of a given clade appear to have lost the paralog. A hatched X shows that a subset of species of the clade have lost it. Filled circles indicate that all species surveyed of the clade retain it. A hatched circle shows that a subset of species of the clade retain it. Open circles show that all species surveyed of the clade appear to have lost it. The teleost phylogeny is based on Betancur-R et al. (2017) and Parey et al. (2023).

Gyoja F, Sato K, Yamashita T, Kusakabe TG. An extensive survey of vertebrate-specific, nonvisual opsins identifies a novel subfamily, Q113-bistable opsin. Genome Biol Evol. 2025;17:evaf032.

Betancur-R R, Wiley EO, Arratia G, Acero A, Bailly N, Miya M, et al. Phylogenetic classification of bony fishes. BMC Evol Biol. 2017;17:162.

Parey E, Louis A, Montfort J, Bouchez O, Roques C, Iampietro C, et al. Genome structures resolve the early diversification of teleost fishes. Science. 2023;379:572–5.

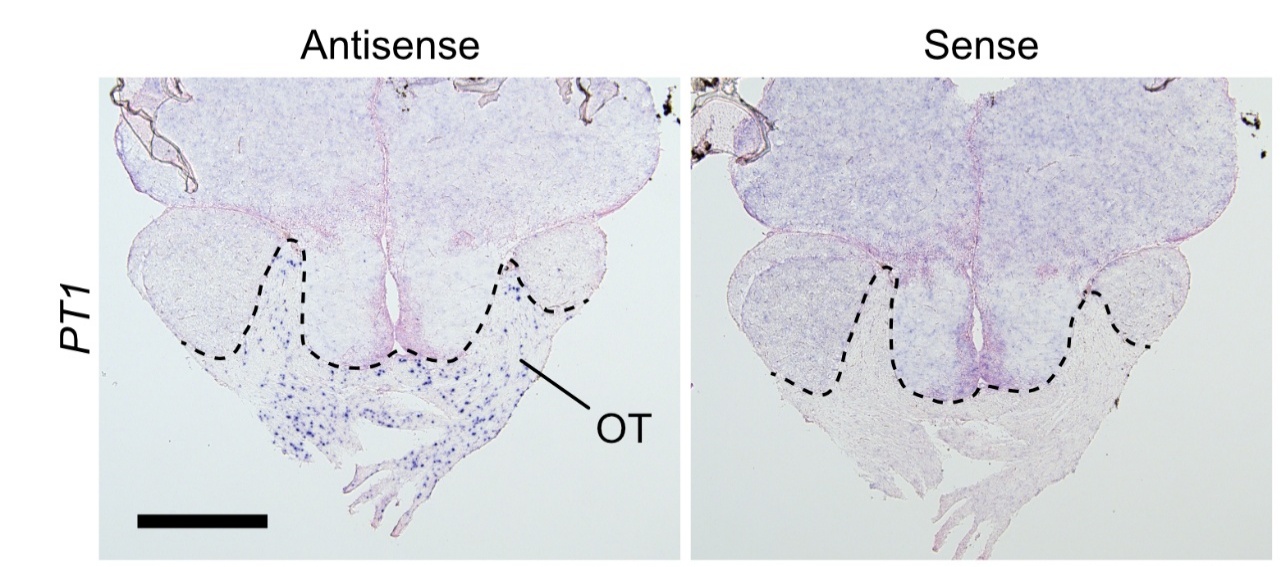
**Supplementary Figure S5**

**Chromogenic *in situ* hybridization for *PT1* in the red piranha brain.**

Consecutive sections hybridized with antisense and sense probes for *PT1*. Dorsal is up. Dashed lines delineate the boundary of the optic tract (OT). Sections were counterstained with Nuclear Fast Red. Scale bar, 500 μm.

**Supplementary Figure S6**

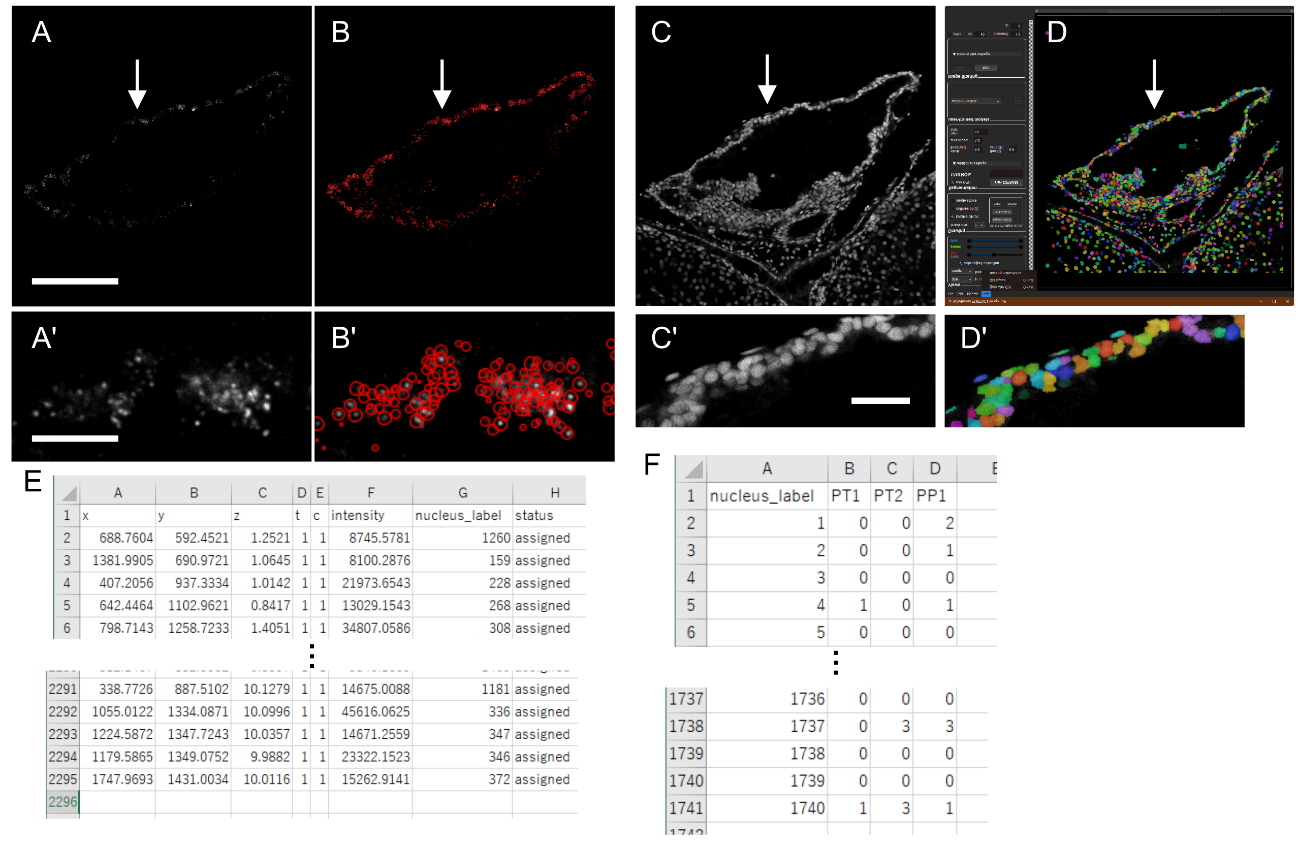

**Workflow for FISH–based cell counting and co-expression analysis.**

(A) Representative *PT2* FISH signal in the red piranha pineal gland (shown as a 2D representation for clarity). (B) RS-FISH spot detection output for *PT2* in the same field of view; detected transcript spots are overlaid (red circles). (C) DAPI nuclear counterstain used for cell identification. (D) Nuclear segmentation result obtained with Cellpose-SAM; nuclei are shown as color-coded instance masks over the DAPI image (top) and as an enlarged overlay (D′). (A′–D′) Magnified views of the regions indicated by arrows in A–D, illustrating spot detection and nuclear segmentation at cellular resolution. (E) Example rows from the RS-FISH spot-localization table exported as a CSV file, including x–y–z coordinates, intensity metrics, the assigned nucleus label (segmentation ID), and assignment status. (F) Example of the per-nucleus summary table (CSV) reporting the number of assigned spots per gene (*PT1*, *PT2*, and *PP1*) for each nucleus label; these per-cell spot counts were used for downstream co-expression analyses (e.g., Venn/overlap statistics) and quantification. Data acquisition, spot detection, nuclear segmentation, and spot-to-nucleus assignment were performed in three dimensions (confocal z-stacks), and spot assignment was based on 3D (x–y–z) distances to the nearest segmented nucleus; panels are shown as 2D representations to facilitate visual interpretation. Scale bars: 100 μm (A; applies to A–C), 10 μm (A′; applies to A′ and B′), 20 μm (C′).

**Supplementary Figure S7**

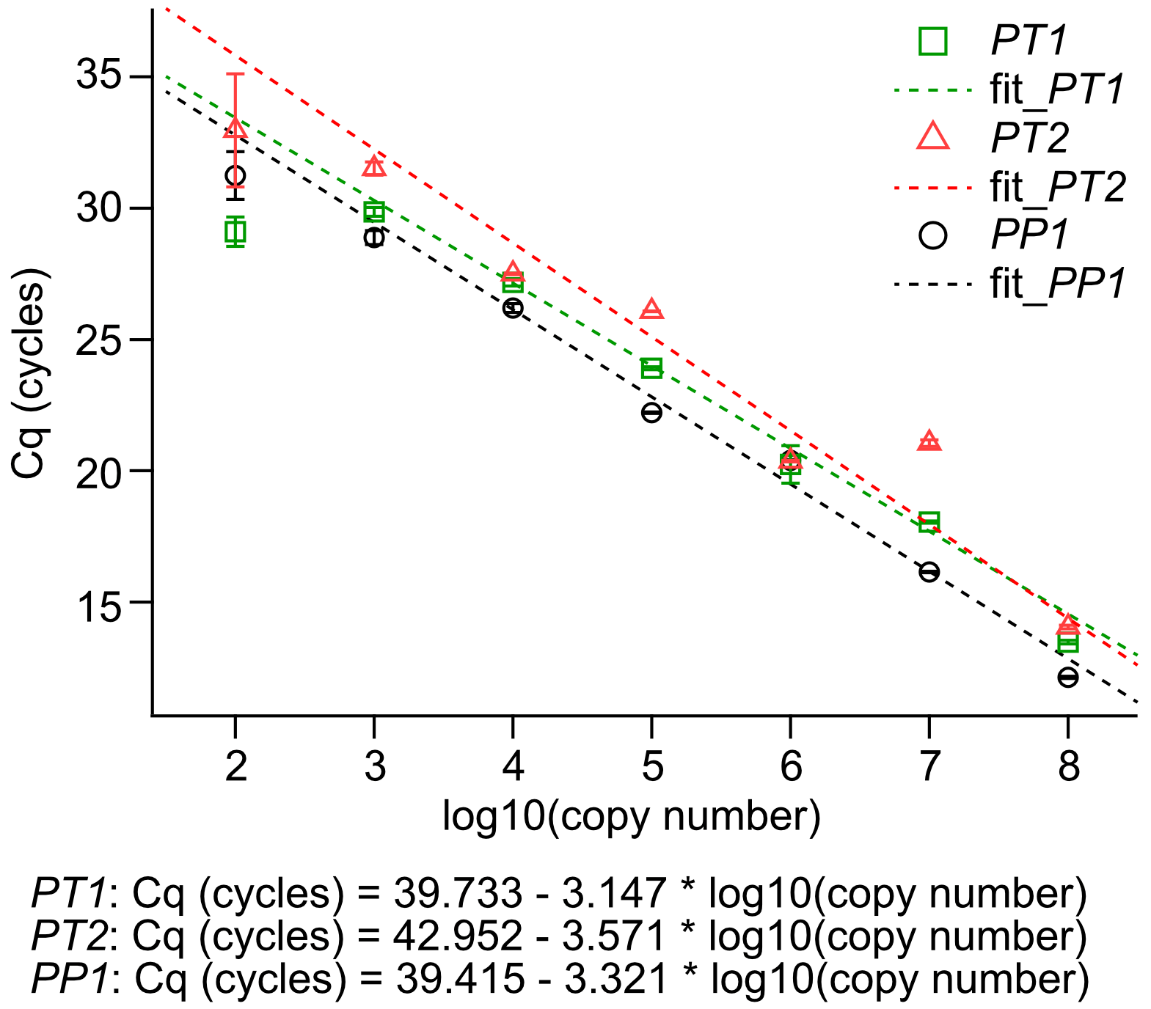

**RT–qPCR standard curves generated from in vitro–transcribed RNA standards.**

Standard curves for *PT1*, *PT2*, and *PP1* were constructed using serial dilutions of synthetic RNA (10²–10⁸ molecules/μL; plotted as log₁₀(copy number)). Following reverse transcription, qPCR was performed and Cq values were plotted against log₁₀(copy number). Symbols indicate measured data points for each target (*PT1*, green squares; *PT2*, red triangles; *PP1*, black circles), and dashed lines indicate the corresponding least-squares linear regressions used for absolute quantification. Regression equations are shown below the plot. Error bars indicate variability among technical replicates (mean ± SD).

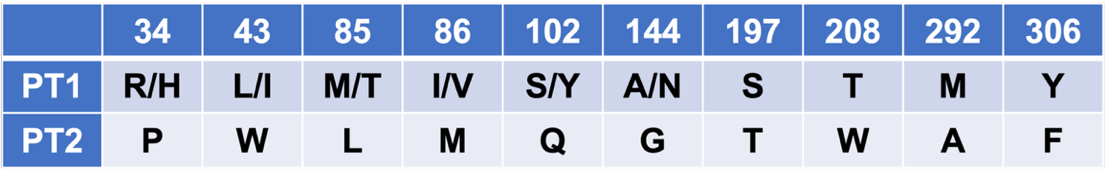
**Supplementary Figure S8**

**Amino acid residues that are conserved within either the PT1 or PT2 group.**

Amino acids are numbered based on the bovine rhodopsin numbering system. See Supplementary Data 1 for the sequence alignment.
